# Fast, Accurate Polarization and Polarity Imaging with Polarized Structured Illumination

**DOI:** 10.1101/640268

**Authors:** Karl Zhanghao, Wenhui Liu, Meiqi Li, Xingye Chen, Chunyan Shan, Haoqian Wang, Peng Xi, Qionghai Dai

**Affiliations:** Department of Biomedical Engineering, College of Engineering, Peking University, Beijing 100871, China; Department of Automation, Tsinghua University, Beijing 100084, China; College of Life Sciences, Peking University, Beijing 100871, China

**Keywords:** optical sectioning, polarization modulation, structured illumination, lipid order, lipid polarity

## Abstract

The orientation and wobbling behavior of the fluorescent dipoles are of great significance in revealing the structure and state of cells. Due to the poor optical sectioning capability of wide-field microscopy, the polarization modulation signals are susceptible to the neighboring fluorophores. The missing cone of wide field optical transfer function induces vast out-of-focus background, resulting in biased polarization orientation and decrease polarization factor. Here, we apply polarized structured illumination to achieve polarization modulation imaging with optical sectioning, and simultaneously measure the lipid polarity with two-color ratiometric imaging. Our results demonstrate a significant increase in measurement accuracy of not only the dipole orientations but also the wobbling behavior of the ensemble dipole. Compared to the conventional confocal polarization imaging, our method obtains an order-of-magnitude faster imaging speed, capturing the fast dynamics of subcellular structures in live cells.

## Introduction

Fluorescence microscopy lights up subcellular structures through specific labeling of biological molecules. The fluorescence has several fundamental physical dimensions including intensity, spectrum, lifetime, and polarization. Fluorescence polarization microscopy measures the dipole orientation of the fluorophore, which indicates the molecular orientation of biological structures^1–4^. The modulation depth defines the degree of the fluorescent signals response to the change of excitation polarization, which relates to the orientational distribution of the dipoles, or the wobbling angle of the ensemble dipole^2,5–8^. It was also previously termed as polarization factor^6,7^, or orientation uniform factor^8^. Measurement of the dipole orientation and polarization factor poses grand importance in the study of biological subcellular structures and interactions^6–12^. However, it can be easily influenced by the out-of-focus background signal in wide-field microscopy, especially for thick specimen. Out-of-focus fluorescence background would lead to decreased modulation depth because the background signal is an average from a large volume of the specimen. Also, a strong polarized background could bias the orientation measurement of the dipoles.

Confocal laser scanning microscopy^12–16^ and spinning disk confocal microscopy^5^ could achieve optical sectioning through the application of confocal pinhole, which were utilized to accurately measure the dipoles with polarization modulation or polarized detection. However, the scanning method in confocal microscopy suffers from slow imaging speed, and polarization modulation in confocal fluorescence microscopy makes the imaging speed even slower. Parallelized acquisition with wide-field imaging can significantly increase the imaging speed, but it can’t remove the influence of out-of-focus background signal on the polarization analysis.

To achieve fast and accurate dipole imaging, we combine optical sectioning^17–22^ wide field imaging with polarization modulation. By introducing polarized structured light^23^, we achieve the polarization modulation while implementing the HiLo sectioning technique. The optical sectioning is achieved by structured illumination for each polarization angle, and polarization analysis is achieved by three or more modulations of the polarization angle of the excitation light. We first imaged the actin filaments in muscle cells, and lipid membrane in U2-OS cells with optical sectioning polarization modulation (OS-PM). For the actin filaments in muscle cells, the polarized out-of-focus background leads to a biased dipole orientation measurement with wide filed polarization modulation (WF-PM), while the OS-PM successfully retrieves unbiased dipole orientation. For the lipid membrane in U2-OS cells labeled with Nile Red, the unpolarized out-of-focus background results in a significantly smaller modulation depth measurement with WF-PM. In contrast, OS-PM can retain the modulation depth, which makes it possible to measure the lipid order of the membrane quantitatively. In addition, OS-PM is capable of capturing the dynamics in live cell imaging with fast imaging speed (30 f.p.s), which has an order-of-magnitude enhancement compared to polarization modulation imaging based on spinning disk confocal microscopy (0.4 f.p.s).

Furthermore, we perform two-color ratiometric imaging of lipid membrane labeled with Nile Red, whose spectrum property indicates the polarity of lipid membrane^24–27^. The intensity ratio from different wavelengths channels reflects the environment polarity in which the fluorophore locates. Similar to the modulation depth, the out-of-focus background also deteriorates the measurement of lipid polarity, for which previous lipid polarity imaging was usually performed with confocal or two-photon microscopy. Two-color OS-PM could image the lipid order for polar and nonpolar lipid membrane in an accurate and fast manner, which is essential for studying the lipid fluidity. The multi-dimensional information of the lipid membrane and its real-time dynamics could provide insights for the biological research in related fields.

## Materials and Methods

### Principle of OS-PM

The original HiLo technique requires one image with uniform illumination and the other with structured illumination. To ensure the simplicity of polarization modulation without switching of the light path, we acquired both images with structured illumination, whose patterns are complimentary to each other. We use *S*(**r**,*α*)to denote the dipole intensity distribution with spatial coordinate **r** and orientation *α*. When the in-focus dipoles are excited by a polarized structured illumination, the number of emitted photons would be *P_i,in_* = *c_in_I_i_S_in_*(**r**,*α*)cos^2^(*θ_i_* – *α*), with *I_i_* = *M_i_* cos(2*π***k_i_ · r** + *φ_i_*)+1. The subscript *i* denotes the index of the illumination pattern, with the pattern direction *θ_i_*, the spatial frequency vector **k_i_**, the phase of structured pattern *φ_i_*, and the modulation contrast *M_i_*. The polarization in polarized structured illumination is parallel to the pattern direction *θ_i_*, which would be explained in the section of optical setup. The dipoles under polarized excitation would be preferentially excited with the quantitative relationship of the cosine square function. The constant coefficient *c_in_* includes the information of the laser power, the quantum yield of the fluorophore, etc. The out-of-focus dipoles are hardly modulated by the structured illumination, and they also have weak polarization modulation due to the ensemble of many dipoles, so that the number of emitted photons from out-of focus dioples would be *P_i,out_* = *c_out_S_out_*(**r**). The polarized out-of-focus ensemble dipole will be rejected by the optical sectioning so it only affect wide field results.

For each pattern direction (excitation polarization), two patterns with exactly a phase shift of *π* were applied, and corresponding emission fluorescence intensity would be Eq. 1 with 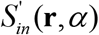 denoting polarization modulated signals at each position 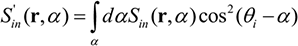. Here the dipoles at the same position are integrated because the camera cannot distinguish them.

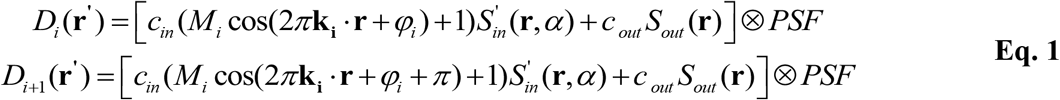

Here the index *i* is an even integer, and the spatial frequency **k_i_**, the modulation contrast *M_i_*, and the polarization 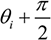 are exactly the same for two patterns. We extract the in-focus dipole information from Eq. 1 with a similar approach utilized in HiLo^22^.

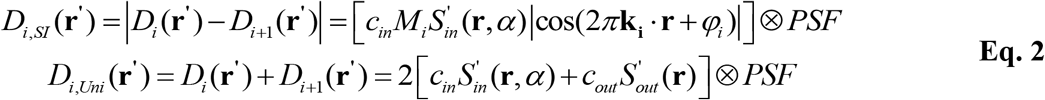

The obtained image modulated with the structured illumination *D_i,SI_*(**r**′) only contains signal from the in-focus dipoles. By applying a low-pass filter (LP) of cutoff frequency smaller than the spatial frequency **k_i_**, we could obtain the low-frequency components of in-focus signals.

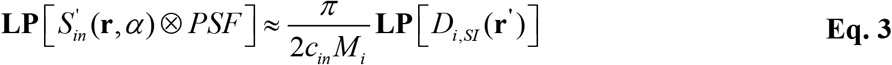

To recover the high-frequency information, we could apply a complementary high-pass filter (HP) to the uniform illuminated image *D_i,Uni_*(**r**′). The out-of-focus background is mostly blurred and only contains low-frequency structure, which means 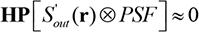. Then we could obtain

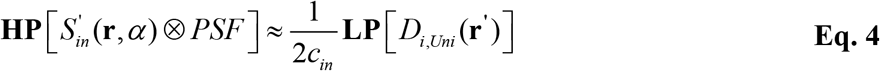

Finally, we obtain a full bandwidth image of the in-focus dipoles with optical sectioning, given by

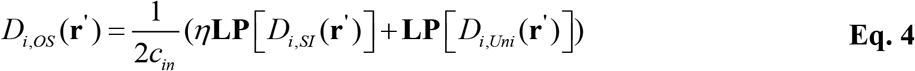

where *η* should be equal to 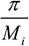 but is treated as an adjustable parameter. One should notice that the modulation contrasts for different pattern direction (polarization) are somewhat different because of imperfect polarization modulation of the illumination laser beam.

Three polarizations are minimal requirements to solve the dipole, while more measurements could increase the precision of measurement under noisy conditions^5^. The specimens in our work behave strong fluorescence, so that we perform three pattern directions in the research. Obtaining the three polarization modulated images with optical sectioning, the dipole orientation could be retrieved by demodulation algorithms. The polarization modulated image with optical sectioning is

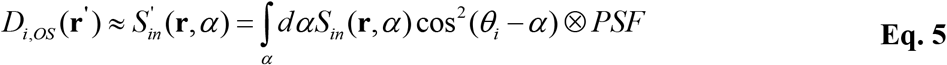

for three pattern directions (i = 1,2,3). The modulation signal of linearly polarized excitation from each pixel is actually from all the dipoles within the diffraction-limited area, which could be taken as an ensemble dipole with average orientation and polarization factor^6–8^. Eq. 5 could be rewritten with

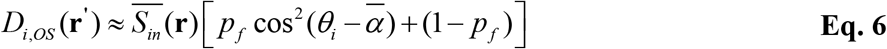

The average orientation 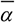, polarization factor *p_f_*, and the intensity 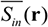 of the ensemble dipole on each pixel could be solved with least-squares estimation describe previously^8^. The term “polarization factor” used here was also called “orientation uniform factor” in the reference, whose definition is the same, as well as the term “modulation depth” used in the paper. “Modulation depth” emphasizes the polarization modulation response of the ensemble dipole, while polarization factor or orientation uniform factor focuses more on the wobbling orientation of the ensemble dipole.

### Optical setup of OS-PM system

The polarized structured illumination is achieved with the system described before^23^. The ferroelectric liquid crystal spatial light modulator (SLM; Forth Dimension Displays, SXGA-3DM-DEV) generates the diffraction pattern. A polarization beam splitter (Thorlabs, CCM1-PBS251) behind the SLM guarantees the laser beams linearly polarized with their polarization parallel to the optical table. Other orders except the ±1 diffraction orders were blocked by a spatial mask. We use a vortex half-wave plate (VHWP, Thorlabs, WPV10L-532) to modulate the polarization of the two ±1 order beams to be parallel to the interference stripes. Two identical dichroic mirrors DM1 and DM2 (Chroma, ZT405/488/561/640rpcv2) were placed perpendicular to each other to compensate for the polarization distortion introduced by them. A 4f system was used to relay ±1 order light spots to the back focal plane of the objective (Nikon, CFI Apochromat TIRF 100× oil, NA 1.49). Two linearly polarized continuous-wave lasers 488-nm (OBIS, Coherent, Santa Clara, CA, USA) and 561-nm (CNI, MGL-FN-561nm-200mW) were used. The emitted fluorescence passes the emission filter (Chroma, ZET405/488/561/640mv2) and a dichroic mirror DM3 (Chroma, ZET635rdc), and finally reach an sCMOS camera (Tuscen, Dhyana 400BSI) to achieve two-color acquisition.

To obtain accurate measurement of the dipoles, the non-uniformity of the laser beams among different pattern directions (polarizations) is calibrated. As the laser beams rotate with the diffractive patterns, any displacement among the center of the Gaussian beams would lead to a variation of excitation power, which tangles with the polarization modulation signal. To calibrate the non-uniformity of the laser beams, we prepared the slide of 100 nm fluorescent beads at a proper density so that the beads can be localized separately. The beads are imaged for two opposite phases for each direction and summed to obtain a wide field image. In each wide field image, the beads are localized by with their position and intensity exported. Those beads appearing on all three images at the same position are used to compensate illumination non-uniformity among different patterns. We either use a quantic polynomial function to fit the non-uniformity or move the beads at a step size of 500 to calibrate the whole field-of-view. Compensation of intensity non-uniformity is performed after optical sectioning by dividing the relative excitation power of each pattern direction.

### Sample Preparation

Human osteosarcoma (U2-OS) cell lines were cultured in Dulbecco’s Modified Eagle’s medium (DMEM, M&C GENE TECHNOLOGY LTD, BEIJING, China) containing with 10% heat-inactivated fetal bovine serum (FBS, GIBCO, USA) and 100 U/ml penicillin and 100 μg/ml streptomycin solution(PS, GIBCO, USA) at 37°C for in an incubator with 95% humidity and 5% CO_2_. The medium was refreshed every three days, and the cells were passaged every week until the cells overspread more than 80% of the petrie dish. 10ng/ml Nile Red (Invitrogen, USA) in the culture medium labels the cells at 37°C for 30-60min.

Muscle cells were acquired from adult mouse and were fixed by 4% formaldehyde (Molecular Probes, USA). Then 1 test of Alexa Fluor 568 phalloidin (Invitrogen, USA) was added to the dish for 10min and then washed three times with PBS. Finally, the slide was sealed with valap (Marabu, Germany) and observed under OS-PM system.

### Data Acquisition and Analysis

At least 6 polarized structured illumination patterns are required to reconstruct an optical sectioning polarization map (as shown in Figure 1(a)). The six patterns can be divided into three groups with polarization modulation orientations being 0°, 60°,120° respectively. The structured illumination of the two patterns in the same group differs only by 90° in phase. The polarized wide field pattern can be obtained by directly superimposing two structured illumination patterns of the same group.

**Figure 1.**
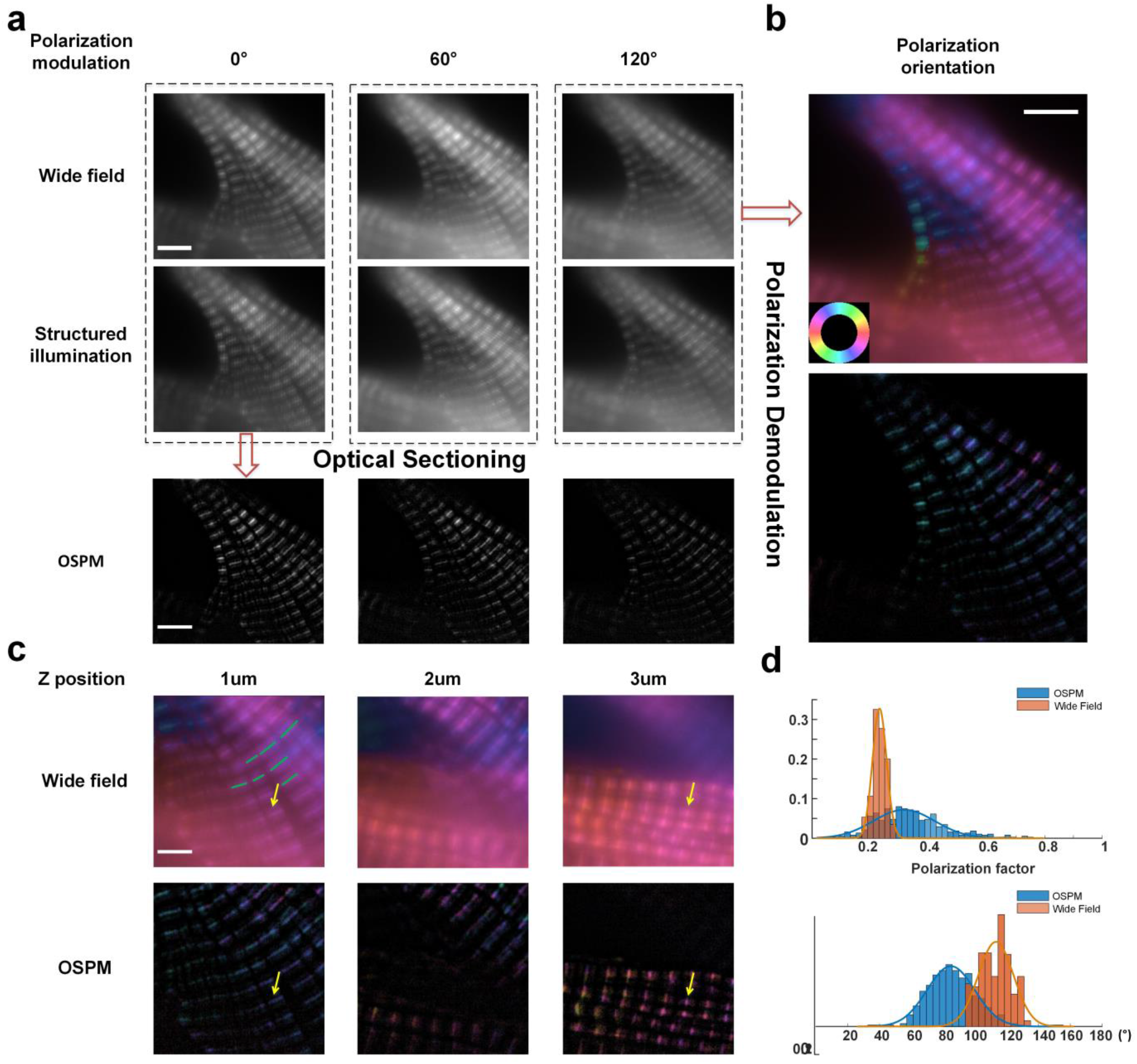
Principle of OS-PM. (a) Wide field images, structured illumination images, and optical section images solved by their combination; each column from left to right represents the structured pattern of 0°, 60°, 120. (b) The dipole orientation mapping images: the figure on top is the wide field result, and the figure below is the OSPM result. The dipole orientation is indicated by the pseudo color the color wheel. For example, red color represents horizontal orientation, and cyan color represents verticle orientation. (c) The polarization analysis of the samples at depths of 1um, 2um, and 3um, respectively. At the position indicated by the yellow arrow, the polarization orientation of the wide field image at 1 um depth is disturbed by the underlying defocused light (structures located at 3 um depth), which is avoided in the OSPM result. (d) Histogram of polarization factor and dipole orientation accuracy by statistics on the position of the green line in figure (c). The dipole orientation accuracy is represented by the angular difference between the polarization direction and the line (M line). The gound truth of angle difference should be 90 degree. Scale bar: 5um in (a) and (b), 3um in (c).

**Figure 2.**
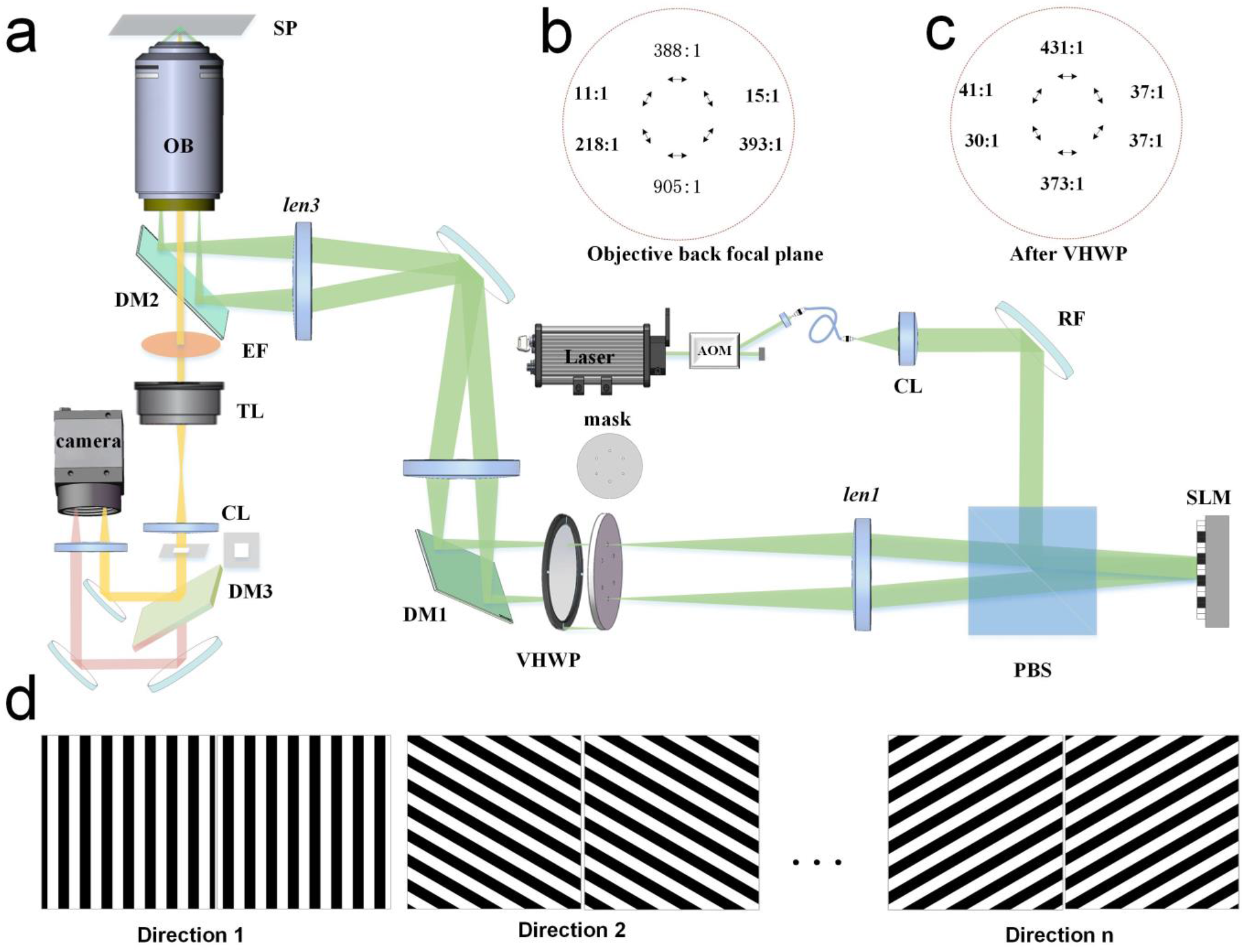
System setup of OS-PM. (a) Schematic setup of OS-PM. AOM: Acoustic Optical Modulator; CL: Collimating Lens; RF: Reflecting Mirror; SLM: Spatial Light Modulator; PBS: Polarization Beam Splitter; VHWP: Vortex Half-wave Plate; DM: dichroic mirror; OB: Objective; SP: Sample Plane; EF: Emission Filter. (b, c) Polarization behavior of OS-PM. (d) Diffractive patterns used in OS-PM.

**Figure 3.**
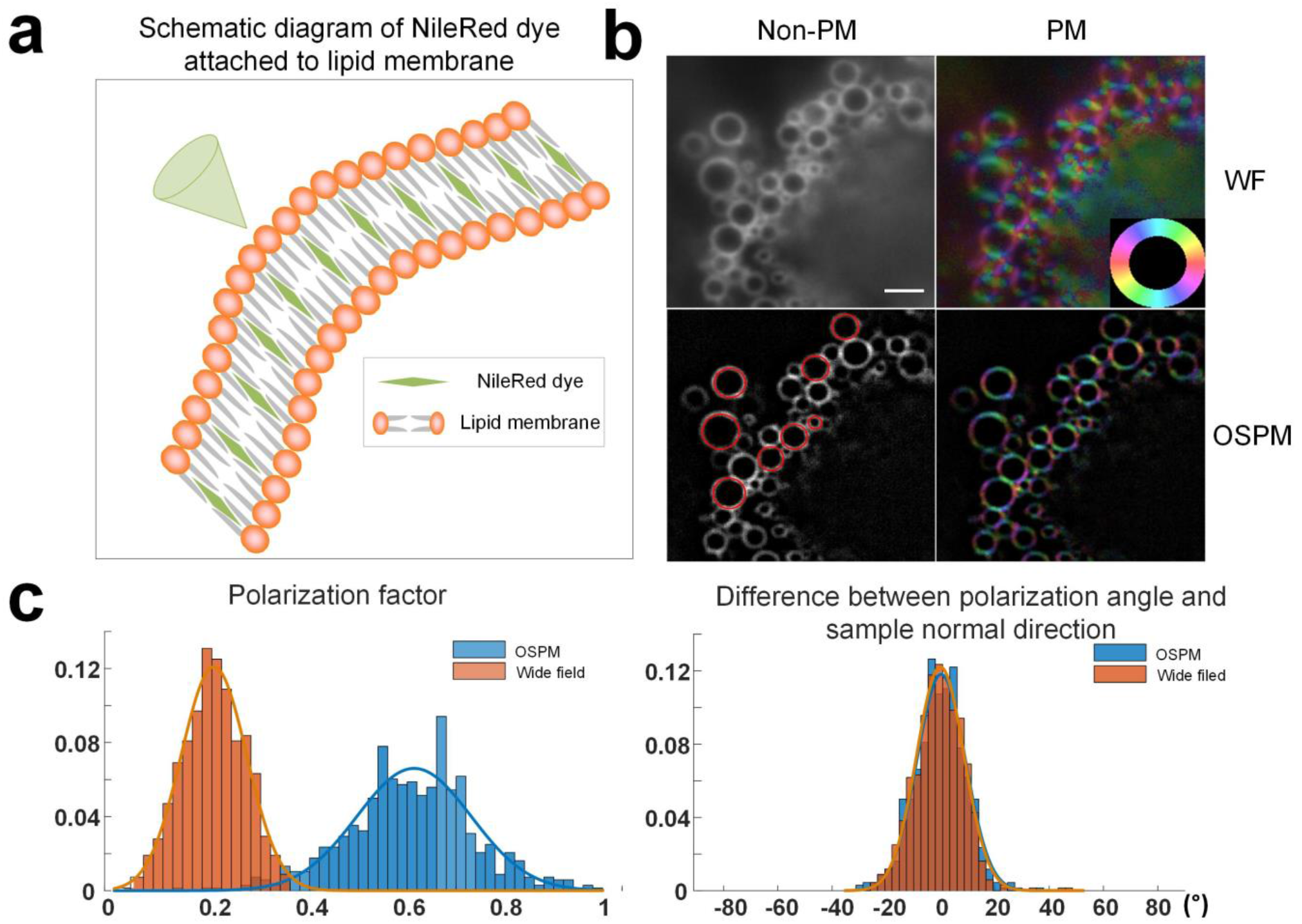
OS-PM increases measurement accuracy of lipid order on the intracellular membrane. (a) The schematic diagram of Nile Red dye attached to lipid membrane, which shows that the direction of the dye molecules is perpendicular to the tangential direction of the lipid membrane. (b) Comparison of the polarization analysis results of wide field and OSPM, the left shows intensity images and the right shows polarization maps. (c) Histogram of polarization factor and dipole orientation accuracy by statistics on the position of the red circle in figure (b). The dipole orientation accuracy is represented by the angular difference between the polarization direction and the sample normal direction. Scale bar: 2um.

The reconstruction process is as previously described. To perform statistical analysis of the performance between WF-PM and OS-PM, we drew the M lines on the muscle cell images by hand and calculated their angular difference between the dipole orientation and the line direction. For the lipid results, we drew the circles on the membranes and calculated their angular difference between the dipole orientation and the normal direction. The distributions are plotted in histograms and fitted with a Gaussian model.

## Results and Discussion

### OS-PM improves the imaging accuracy of the dipole orientation

The polarized out-of-focus background would bias the measurement of the in-plane ensemble dipole towards the out-of-focus polarization. With optical sectioning rejecting the out-of-focus background, the measurement accuracy of the dipole orientation could be significantly improved. To verify our idea, we prepared the sample of muscle cells labeled with phalloidin conjugated Alexa Flour 568. Previous studies showed that the dipole of AF568 fluorophores would orient along the actin filament^23^. The actin filaments in the muscle cells are well known with organized distribution perpendicular to the M lines which was indicated by the lines in Figure 1c. We imaged an area with multiple muscle cells overlaid with a different orientation. A z-stack of the actin filaments in Figure 1c clearly shows different orientations of the actin bundles exist in a different focus. When the in-focus dipoles (z=0 um) are imaged, their polarization responses tangled with the defocused dipoles (z= 3 um) with the orientation of 156.5°. We analyzed the orientation of the ensemble dipole on the lines in Figure 1c, whose ground truth would be perpendicular to the lines (about 117.6°). Conventional polarization modulation based on wide-field imaging results in an average orientation of 157.3°, which is biased by the defocused polarization background. The OS-PM result leads to an average orientation of 116.4°, which is in better accordance with the ground truth.

### OS-PM improves the imaging accuracy of the polarization factor

Because the blurred background contains polarization response from a large number of dipoles, the out-of-focus signal is weakly polarized or unpolarized in many cases, which will not affect the measurement of the dipole orientation *α*. However, the unpolarized out-of-focus background would reduce the depth of the polarization modulation, resulting in a reduced polarization factor. A smaller polarization factor means a larger wobbling orientation of the ensemble dipole, or a more disorder distribution of the dipoles. This approach has been widely used in previous papers to indicate the lipid order of the membrane^5,13,15,28^. We imaged the lipid order on the intracellular membrane with Nile Red labeling. Since Nile Red binds to a large variety of lipid membrane^26^, the out-of-focus background is severe. The dipole of Nile Red inserts into the membrane perpendicularly, which could be measured accurately by both conventional PM and OS-PM. Nevertheless, the polarization factor measured by OS-PM has a much larger value (with an average of 0.61) than the conventional PM method (with an average of 0.20). The OS-PM result rejects the constant unpolarized out-of-focus background, which is more reliable.

### Two color OS-PM simultaneous measures the lipid order and polarity

For lipid membrane staining dye of Nile Red, its emission spectra are influenced by the polarity of the environment^25,27^. Nile Red in nonpolar environment such as lipid droplet emits mostly green-yellow fluorescence, while Nile Red in polar environment emits more red fluorescence. Therefore, the polarity of the membrane could be obtained using ratiometric imaging. We spilt the emitted fluorescence of Nile Red onto two halves of the sCMOS camera.

The excitation wavelength is 561 nm, and a highpass dichroic mirror (635 nm) is placed before the detector. The ratios between the two channels reflect the polarity of the lipid membrane. The accuracy of measured spectral ratio would also benefit from optical sectioning, because the out-of-focus background would add values similarly to both channels, making the spectral ratio close to 1. Furthermore, we performed OS-PM on each channel so that we could obtain the lipid order of both nonpolar membrane and polar membrane. The statistical results show a smaller polarization factor in nonpolar enviroment than in polar enviroment in Figure 4(e).

**Figure 4.**
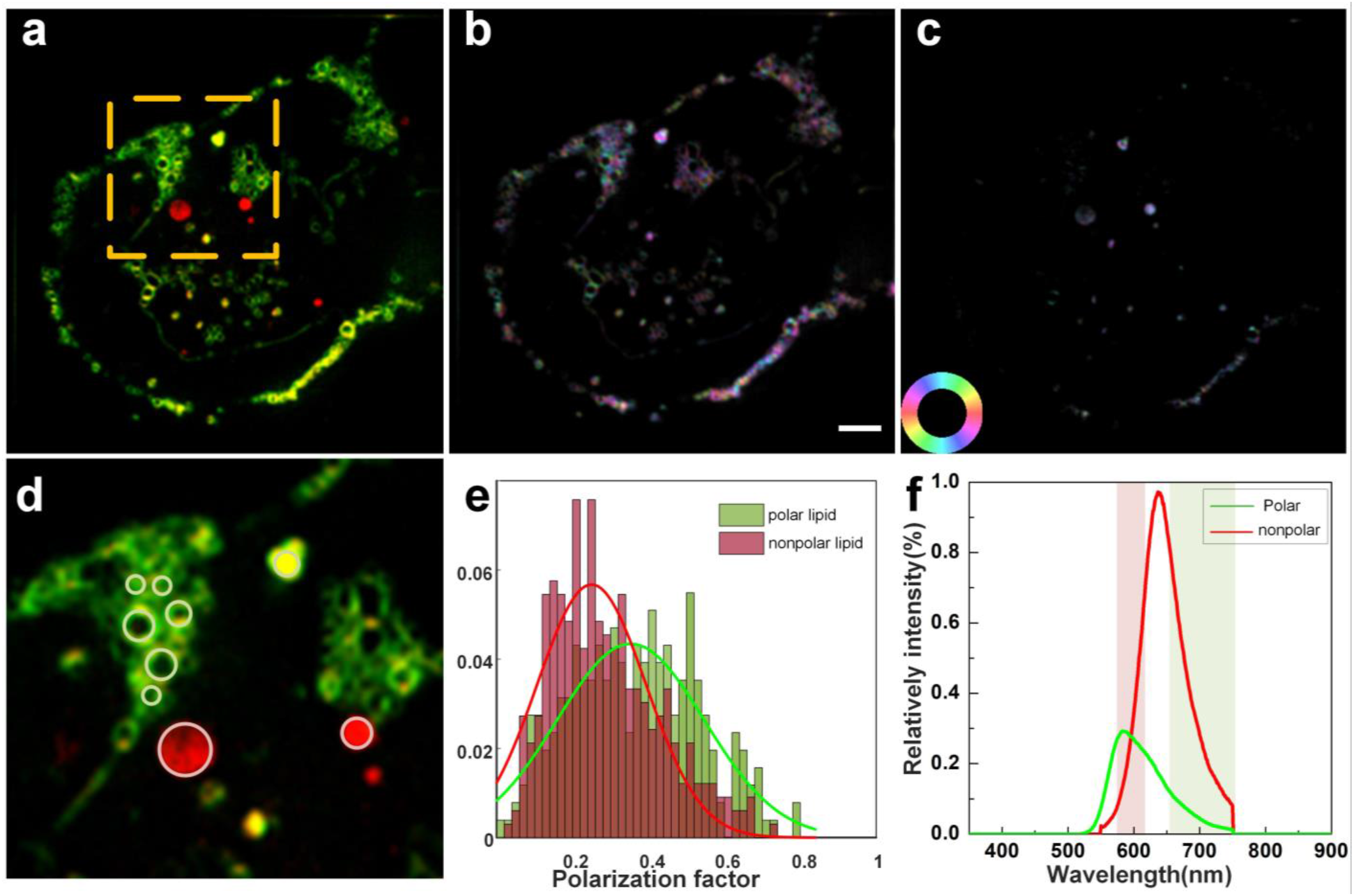
OS-PM increases measurement accuracy of lipid order on the intracellular membrane. (a) Dual color merge image of lipid membrane labeled with Nile red while red indicates nonpolar lipid and green denotes polar lipid. (b) Polarization mapping image of lipid membrane in polar environment. (c) Polarization mapping image of lipid membrane in nonpolar environment. (d) The zoom-in image of (a). (e) Statistical histogram of polarization factor at the position of circles in figure (d). The red histogram shows the statistical results of nonpolar lipid circled in pink while green histogram illustrated the results of polar lipid circled in reseda in (d). (f) The emission spectrum of two different components of lipid at 561 nm excitation. Scale bar: 3um.

**Figure 5.**
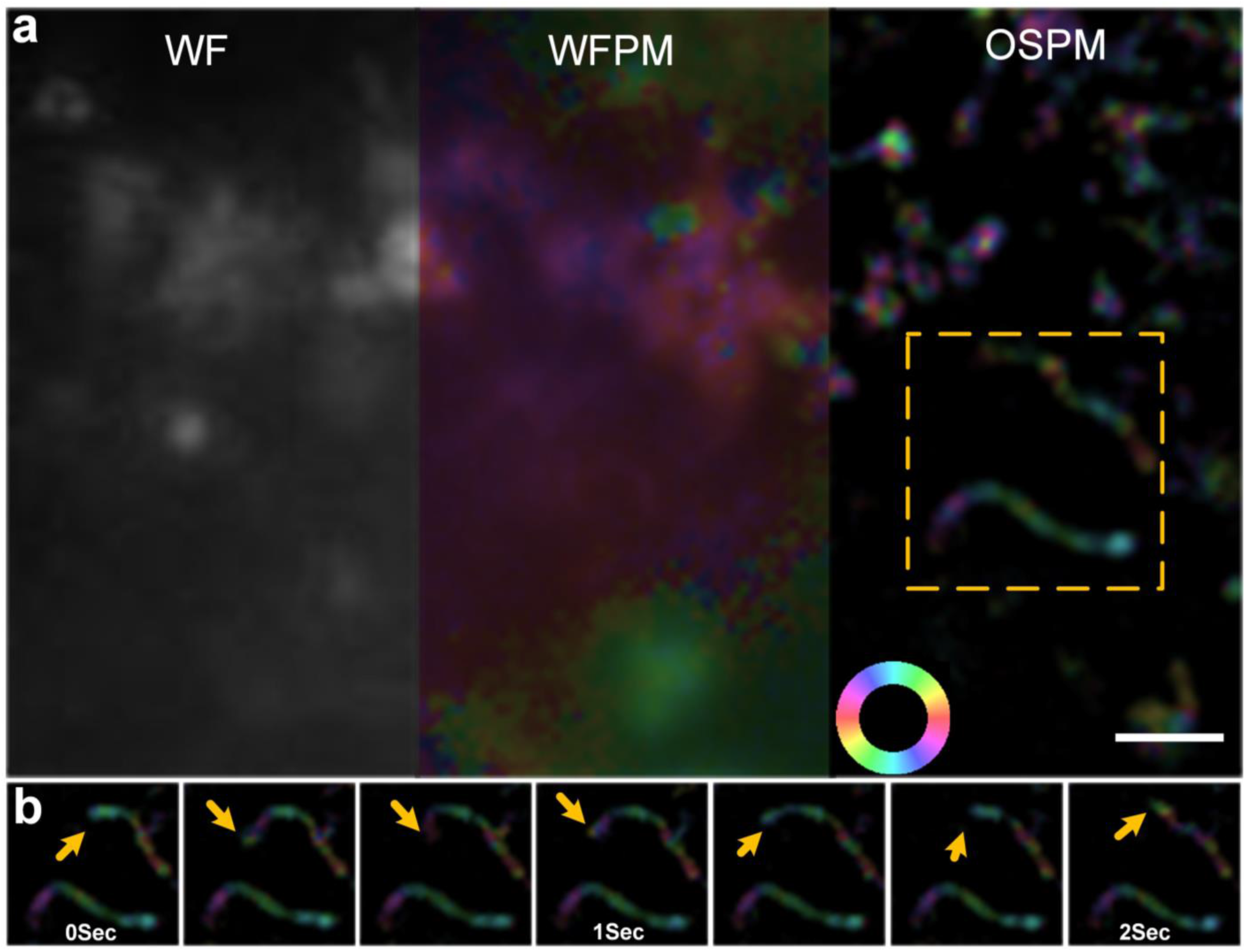
OS-PM tracks fast dynamics of lipid membrane in live cells. (a) Imaging results of different subcellular structure labeled with Nile red at fast speed 30 f.p.s. Wide field (left), wide field polarization modulation (middle) and optical sectioning polarization modulation (right). (b) The zoom-in image in (a) and the time-lapse images. Scale bar: 2um.

### OS-PM captures fast dipole dynamic in live cells

Polarization modulation based on confocal or spinning disk confocal could also obtain optical sectioning and achieves accurate imaging of the dipole orientation and polarization factor. However, the point scanning methods have the drawback of slow imaging speed. Polarization modulation would make the acquisition time even several times longer. In contrast, OS-PM could achieve fast imaging speed with parallelized acquisition of a digital camera. The display speed of the diffractive pattern achieved by the SLM can be as fast as 3.2KHz, and the imaging speed of our sCMOS camera is able to achieve hundreds of frames with an ROI of 512*256 pixels. Therefore, the limiting parameter of the imaging speed is ideed the signal-to-noise ratio of the fluorophore. With an exposure time of 5 ms, we could acquire 186 frames per second, which leads to the imaging speed of 31 reconstructed frames per second. The fast imaging speed of OS-PM has an order-of-magnitude improvement compared to confocal based polarization modulation. Fast imaging speed is curial for the measurement of dipole orientation, because the motion blur of dipoles with a polarization modulation cycle would induce additional intensity fluctuation, which could bias the measured dipole orientation. To demonstrate the capability of OS-PM on fast live cell imaging, we performed time-lapse imaging of the Nile Red labeled lipid membraned in live U2-OS cells. The dipole orientations measured on fast moving lipid membranes are not disrupted by the motion blur and are still perpendicular to the membrane.

### Limitation of our works

There are still several limitations to our OS-PM technique. Firstly, the excitation polarization remains within the sample plane in OS-PM, which only measures the 2D dipole orientation of the fluorophore. In most cases, the dipole has a 3D-distribution of orientation and wobbling behavior. The out-of-plane tilt angle of the 3D dipole will not affect the measurement accuracy of the in-plane orientation, while it leads to a biased measurement in wobbling angle of the ensemble dipole^5^. Therefore, we have to select the specific sample plane in which the dipoles lie in the focal plane. Secondly, similar to confocal or spinning disk, optical sectioning based on HiLo principle rejects a fraction of in-focus signal at the same with rejecting the out-of-focus background. Therefore, the signal-to-noise ratio of the images is reduced, which also limits the imaging speed of OS-PM. A better approach is to separate all the in-focus signals from the out-of-focus background, which may require a 3D structured illumination pattern. Thirdly, though two color OS-PM improves the measurement accuracy of the spectral ratio of Nile Red, a quantitative study of the lipid polarity requires calibration experiments under the same imaging condition with manufactured lipids with different polarities, which remains to be our future work. Lastly, even though the excitation lasers are close to linearly polarization, the remaining ellipticity could still be included in the demodulation model to further improve the accuracy of measured dipoles^15^.

## Conclusion

To sum up, we achieved optical sectioning polarization modulation with six images under polarized structured illumination. While wide field based polarization modulation suffers from the out-of-focus background, OS-PM significantly improves the measurement accuracy of dipole orientation and polarization factor. Compared to confocal based polarization modulation, OS achieves an order-of-magnitude improvement of the imaging speed (30 f.p.s.). We demonstrated the accurate measurement of dipole orientation of OS-PM on the actin filaments in muscle cells, while the WF-PM measurements are biased by the polarized out-of-focus background. Afterwards, OS-PM accurately measures the polarization factor of the Nile Red dipole, which reflects the order and disorder of the lipid membrane. With two color OS-PM imaging, we can simultaneous image the lipid order and polarity and capture their dynamics in live cells. The comprehensive and multi-dimensional measurement of the lipid membrane could hopefully advance the biological researches in related fields.

